# Multiple, single trait GWAS and supervised machine learning reveal the genetic architecture of *Fraxinus excelsior* tolerance to ash dieback in Europe

**DOI:** 10.1101/2023.12.11.570802

**Authors:** JM Doonan, KB Budde, C Kosawang, A Lobo, R Verbylaite, JC Brealey, MD Martin, A Pliūra, K Thomas, H Konrad, S Seegmüller, M Liziniewicz, M Cleary, M Nemesio-Gorriz, B Fussi, T Kirisits, MTP Gilbert, MTP Heuertz, ED Kjær, LR Nielsen

## Abstract

Common ash (*Fraxinus excelsior*) is under intensive attack from the invasive alien pathogenic fungus *Hymenoscyphus fraxineus*, causing ash dieback at epidemic levels throughout Europe. Previous studies have found significant genetic variation among clones in ash dieback susceptibility and that host phenology, such as autumn yellowing, is correlated with susceptibility of ash trees to *H. fraxineus*; however, the genomic basis of ash dieback tolerance in *F. excelsior* remains poorly understood. Here, we integrate quantitative genetics and genome-wide association analyses with machine learning to reveal the genetic architecture of ash dieback tolerance and its relationship to phenological traits in *F. excelsior* populations in six European countries (Austria, Denmark, Germany, Ireland, Lithuania, Sweden). We use whole-genome sequencing of 486 *F. excelsior* genotypes to confirm the genotypic correlation between crown damage caused by ash dieback and intensity of autumn leaf yellowing within multiple sampling sites. Although, our results suggest that the examined traits are polygenic, a relatively small number of single nucleotide polymorphisms (SNPs) explained a large proportion of the variation in both disease tolerance and autumn leaf yellowing. We could explain up to 63% (based on 9155 unlinked SNPs) of variation in individual response to ash dieback crown damage and up to 72% (based on 3740 unlinked SNPs) of variation in autumn yellowing. We identified eight SNPs encoding non-synonymous substitutions, of which those with the highest predictive power were located within genes related to plant defence (pattern triggered immunity, pathogen detection) and phenology (regulation of flowering and seed maturation, auxin transport). Overall, our results provide insights of a multifaceted defence response, according to which a combination of direct defence mechanisms and phenological avoidance of pathogen spread constitute tolerance to ash dieback.

## Introduction

The threat to indigenous plants from new and emerging forest pests and pathogens (PnPs) has substantially increased in recent decades due to the increase in international trade and travel. Globalisation has enhanced trade between distant parts of the planet with concomitant exposure of native plants to introduced pathogens, to which they have no evolved immunity. This has led to devastating and ongoing PnP outbreaks such as *Ophiostoma ulmi* and *Ophiostoma novo-ulmi* causing Dutch elm disease since the start of last century (Brasier, 1991), to devastation of ash trees caused by the translocated Emerald Ash Borer (*Agrilus planipennis*) from East Asia to North America possibly from as early as the 1980s (McCullough, 2020), to chestnut blight caused by the early 20^th^ century introduction of the fungus *Cryphonectria parasitica* from East Asia to North America, causing the extirpation of the once common American chestnut (*Castanea dentata*) (Anagnostakis, 1987). The introduction of PnPs brings novel and strong selection factors to indigenous plants, which in the absence of a shared evolutionary history must solely rely on standing genetic variation to survive and adapt (Budde *et al*., 2016).

Trees have long generation times (in contrast to their PnPs), where polygenic mechanisms of disease resistance offer a broad response spectrum to the cornucopia of threats encountered by a tree during its long life (Bruns *et al*., 2015; Yeaman, 2022). In forest trees, resistance based on a major gene is therefore likely to be rare (but see the case of western white pine trees against blister rust caused by the introduced fungal pathogen *Cronartium ribicola* in North America (Kinloch *et al*., 1999)). Polygenic resistance is not only more frequent in nature but offers a more durable form of resistance, which to be defeated requires multiple virulence genes from the PnP (Bell, 1982; Maher, 2008; Palloix *et al*., 2009).

The development of high-throughput sequencing technology in combination with improved analytical tools facilitate research on polygenic genetic mechanisms underlying forest tree PnP resistance. Polygenic traits where multiple loci contribute with small effects to the phenotype, complicate the interpretation of genotype-phenotype interactions. This is particularly true for Genome Wide Association Studies (GWAS), which have had great success in deciphering the genetic architecture of monogenic traits (Sánchez-Vallet *et al*., 2018), but have struggled to unravel the effects of multiple genes in polygenic traits. However, knowledge of the genetic architecture of polygenic disease resistance is crucial to support the stability of future forests in the face of ever growing threats from PnPs (Stocks *et al*., 2019; Elfstrand *et al*., 2020).

Ash dieback (ADB), a disease of common ash (*Fraxinus excelsior* L) is caused by the invasive ascomycete, *Hymenoscyphus fraxineus* (syn. *H. pseudoalbidus*; anamorph *Chalara fraxinea*) (Kowalski, 2006; Queloz et al., 2011; Baral et al., 2014). The disease was first observed in north-eastern Poland in 1992, before spreading through the Baltics to Denmark in 2002, France in 2008, and the UK in 2012 (Marçais *et al*., 2022). Native to East Asia, *H. fraxineus* evolved as a mild leaf and shoot pathogen of indigenous Asian ash species with insignificant overall impact on the host (Zhao *et al*., 2013; Drenkhan *et al*., 2017). Translocation of *H. fraxineus* to Europe, onto a naïve and compatible host (*F. excelsior*) has initiated the ADB epidemic and resulted in an emerging disease caused by a highly virulent pathogen. Spreading from eastern to western Europe, the range of *H. fraxineus* has no spatial limitations and is only restricted by higher temperatures in southern Europe and the occurrence of ash species (Enderle *et al*., 2019). The disease similarly affects narrow-leaved ash, *Fraxinus angustifolia* but crucially *F. angustifolia* has a more southern European distribution where the pathogen is limited (Dal Maso, 2014). Disease pressure builds in annual cycles where the wind-dispersed ascospores of the pathogen spread to new host trees and re-infect them during the summer months (Marçais *et al*., 2022). Mycelia of *H. fraxineus* spread from infected leaves into woody parts of the tree causing progressive necrosis and crown dieback, which often lead to mortality of trees of all ages. In late summer and autumn, the infected leaves fall to the ground as leaf litter acting as conduits for sexual recombination, the formation of apothecia and the dispersal of new ascospores.

The impact of ADB on *F. excelsior* is, in addition to tree genotype dependent on multiple factors including general tree health, age, environmental conditions, infection pressure, presence of other pathogens and endophytes as well as density and spatial heterogeneity of host populations (Madsen *et al*., 2021; Marçais *et al*., 2022). Previous studies have revealed a polygenic genetic architecture with moderate to high heritability in ADB tolerance among individual trees. These studies revealed narrow sense heritability for ADB (Pliura *et al*., 2011; Kjær *et al*., 2012; Lobo *et al*., 2014; Muñoz *et al*., 2016) of 0.37 to 0.53 and broad sense heritability (McKinney *et al*., 2011; Stener, 2013; Enderle *et al*., 2015) of 0.1 to 0.57 among different ash trials in Europe. Furthermore, the level of crown damage due to ADB is genetically significantly correlated to autumn leaf colouring and almost significantly correlated to spring bud burst (McKinney *et al*., 2011; Stener, 2013). Therefore, it has been speculated that the observed genetic resistance may be partly attributed to differences in the timing of phenological stages. Phenological avoidance of severe disease has been identified in other pathosystems such as Dutch elm disease where the susceptibility of elm trees is related to spring phenology (Ghelardini & Santini, 2009). However, in previous tests of necrosis formation after direct stem inoculations of *H. fraxineus* onto ash clones and progeny, trees that performed better in the ADB-affected trials also had reduced necrosis, which indicates defence-mediated tolerance to ADB (McKinney *et al*., 2012; Lobo *et al*., 2015). The relative contribution of defence mediated tolerance and phenological avoidance to overall ash dieback tolerance is unknown.

A recent study based on high-throughput sequencing and GWAS on ash trees covering the U.K., Ireland and Germany aimed to identify molecular markers underpinning ADB resistance (Stocks *et al*., 2019). This study found polygenic resistance to ADB and association between 61 highly significant SNPs and a binary trait.

Here, we investigated the genetic architecture of *F. excelsior* tolerance to ADB in six countries across Europe, taking advantage of clonal common garden and progeny trials. The aim of the study is to identify associations between the level of crown damage (a continuous trait, which is a proxy for the overall impact of ADB), spring and autumn leaf phenological traits, and molecular markers. Due to the relatively large heritability estimates identified from previous studies and correlations between ADB and phenology, we hypothesise that; 1) genetic variants (SNPs) can explain ash dieback crown damage and phenological variance among individuals. 2) SNPs of importance are located within coding regions and in functional areas related to tree defence and phenology, 3) there are overlaps between SNPs associated with ADB crown damage and SNPs associated with phenology traits. To address the above hypotheses, multiple phenotypic traits were recorded on common ash trees on a quantitative scale to reflect the phenotypic spectrum of ADB symptoms on individual trees and of spring and autumn phenology. Furthermore, GWAS analysis was combined with machine learning to efficiently identify the association between phenotypic observations, namely ADB crown damage, spring bud burst, autumn leaf yellowing, autumn leaf loss and overall autumn status, to underlying molecular markers. Except for the inclusion of a single progeny trial, ash tree leaves were sampled within clonal trials where multiple ramets of each clone were grown across each site, providing accurate quantitative phenotypes, and therefore providing the best opportunity to capture associations between phenotype and genome.

## Material and Methods

### Field assessments and sampling

A set of clonal trials (some originally intended for seed collection) with non-overlapping clones located in Austria, Denmark, Germany, Ireland, Lithuania, Sweden were included in the study (Supporting Information Table S1). All sites were infected by wind dispersed ascospores of *H. fraxineus*, and most trees showed ADB symptoms. The field trials had been monitored for ADB symptoms prior to the present study and significant variation in disease symptoms had been observed among clones (see references in Supporting Information Table S1). Genotypes were selected to include trees in a gradient from high to low crown damage scores. Trees with no or limited ADB crown damage were, however, under-represented in the clonal trials, and therefore a Danish progeny trial (located in Silkeborg) was included in the study.

All trees at each trial were assessed for five traits: spring phenology (i.e. spring bud burst), crown damage with typical ADB symptoms, and three scores related to autumn senescence. The traits were scored simultaneously in all sites (Supporting Information Table S1).

Spring bud burst: The leaf phenological stage of each tree was assessed and scored on a scale from 0-7:

Score 0: bud still in winter stage, 1: Bud swollen, black and/or green bud scales enclose the leaves completely, 2: Buds are beginning to burst and leaves only just visible, 3: Leaves visible, very small and have just escaped the bud, 4: Leaves are unfolding, glossy and the shoot starting to stretch, 5: Leaves stretched markedly, still glossy, 6: Leaves full size, still “spring fresh” and not completely hardened, 7: Mature, all leaves are dim and hardened. Please see Supporting Information Fig. S1 for illustrations. Each tree was given one score that reflected its average spring phenological stage across the entire crown on the assessment day. Several of the sites were scored twice (Supporting Information Table S1) to assure capturing a time point with maximal phenotypic variation among clones.

#### ADB crown damage

trials with young trees were scored on a scale with five categories: 0: No visible symptoms, 1: < 10% of the crown with ADB symptoms, marginal damage on stem and crown, 2: 10-50% of the crown with ADB symptoms, presence of dead parts in the crown and discoloured necrotic parts on stem, 3: > 50% of the crown with ADB symptoms, prominent to highly dominant presence of damage on shoots, branches and stem, including larger necrotic areas on stem and branches, 4: tree dead because of severe infection. For old trees three additional categories were included (see Supporting Information Table S2). To rank clones across the sites, both old and young crown damage scales were transformed into percentage damage score (PDS) using the following five class means (0%, 5%, 30%, 75%, 100%).

#### Autumn leaf yellowing

this trait was scored on a five-step scale reflecting the overall impression of the autumn colour of each tree using the following scale: 0: leaves still dark green, 1: leaves slightly lighter, or dark green but with yellowing leaf nerves, 2: leaves green but with yellow spots on leaflets, 3: yellowing leaflets, 4: completely yellow leaves. See Supporting Information Table S2 and Fig. S2 for detailed descriptions and illustrations.

#### Autumn leaf loss

this trait was scored as percentage of leaves lost in the crown apparently due to autumn senescence and scored in 10% classes (Supporting Information Table S2 and Fig. S2).

#### Autumn status

this trait reflects the overall progress towards dormancy status of a tree; the remaining leaves showing senescence symptoms combined with the progression of senescence symptoms. Symptoms of senescence of foliage: summer green colour becomes lighter, yellowing, brownish, withering (typically from the rim towards the middle) and crusted, dried out or hanging leaves. Thus, the score is gradual and shows average tree senescence level (full dormancy = 100%) (Supporting Information Table S2 and Fig. S1).

A subset of *F. excelsior* individuals which had been phenotypically assessed, were sampled from each site for DNA extraction. Based on data of ADB assessments from previous years we aimed to span the entire ADB crown damage range from each trial when selecting the trees. From each sampled tree, 2-4 fresh leaflets were collected and placed in ziplock-bags with silica gel. The samples were stored at 4°C until further processing.

### Statistical Analyses

Phenotypic data for the GWAS was derived from the ADB crown damage (PDS) and phenology scores described above. Clone least square mean values (LSMeans) were estimated per trial using the model given below using PROC GLM in SAS 9.4:

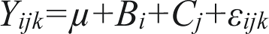

where Y*ijk* is the phenotypic score measured for trees, *µ* is the overall mean of the trial score, *Bi* is the fixed effect of block within a trial, *Cj* is the random effect of clones, and *εijk* is the residual.

Clones were ranked across sites for each phenotypic trait based on the values obtained by subtracting their LSMeans values from the site average for each trait.

Genotypic correlations (rG) among traits were calculated using bivariate analysis in ASReml v4.2 using the following equation:

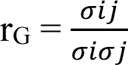

where σij is the clone covariance component between traits i and j, σi and σj are the standard deviations for clone variance components for traits i and j respectively.

### DNA extraction

DNA was extracted from 20 mg of dried leaf tissue using the NucleoSpin Plant II kit (Macherey-Nagel, Denmark). DNA was extracted according to the manufacturer’s protocol with the following modifications: 100 μl of lysis buffer PL1 was added to each sample, an additional initial physical disruption step using a Qiagen Tissue Lyser II (Qiagen, Germany) at 25 rpm/s for 1 min with two 3 mm steel beads for each sample, then 500 μl additional lysis buffer PL2 was added to each sample, prior to two additional physical disruption steps at 25 rpm/s for 1 minute. Subsequently, 10 μl RNase A was added to each tube, the samples were vortexed briefly and incubated at 65°C for 15 minutes and inverted 2-3 times during the incubation. Samples were placed in a centrifuge at 8,000 rpm for 5 mins to remove crude lysate. Supernatant was transferred to a violet ringed NucleoSpin ii tube, and from this point extracted according to the manufacturer’s instructions. Extracted DNA was eluted in 50 μl volumes and concentration was measured using a Qubit 3.0 fluorometer (Invitrogen, USA) and Qubit dsDNA BR kit (Invitrogen, USA).

### Microsatellite analysis

Prior to library preparation two ramets per clone within each site, were genotyped using microsatellite markers to omit potential grafting or sampling errors in the data. In cases of mismatch between two genotyped ramets, the clone was omitted from the study. Microsatellite analyses were performed with three selected primer pairs multiplexed in a mix: FEMSATL11 and FEMSATL19, as well as FEMSATL12 (Lefort & Douglas, 1999; Gerard *et al*., 2006). PCR amplifications were carried out using the Qiagen Multiplex PCR Kit (Qiagen, Germany) according to the manufacturer’s instructions but scaled down to 10 μl reaction volume. PCR amplifications were performed under the following conditions: initial denaturation at 95 °C for 15 min, 30 cycles of denaturation at 94 °C for 30 s, annealing at 57 °C extension at 72 °C for 60 s, and final extension at 60 °C for 30 min. PCR products were analysed on the ABI 3130xl Genetic Analyser (Applied Biosystems, Foster City, CA, USA).

### Library preparation

DNA was sheared using a LE220-plus Focused-ultrasonicator (Covaris, USA), with a target insert length of 500 bp. Sheared DNA fragment size and concentration was assessed using a Fragment Analyzer 5300 with 48 capillaries. The Fragment Analyzer was operated by the National High-Throughput DNA Sequencing Centre (University of Copenhagen, Denmark). DNA libraries were built using the BEST protocol (Carøe *et al*., 2018) with an input DNA quantity of 100 ng. Prepared libraries were PCR-indexed (Supporting Information Table S3) using AmpliTaq Gold polymerase with the following conditions: 95°C for 10 mins followed by 12-16 cycles at 95°C for 20 s, 60°C for 30 s, 72°C for 40s, and a final elongation step at 72°C for 7 mins. DNA Libraries were dual indexed and subsequently pooled into equimolar concentrations with a maximum of 96 samples in each pool.

Pooled DNA was sequenced by Macrogen Europe (Amsterdam, The Netherlands) on the NovaSeq 6000 platform with 150 bp PE reads, using the S4 flow cell workflow. A further 10 samples were sequenced on the Illumina HiSeq 4000 platform at the National High-Throughput DNA Sequencing Centre (University of Copenhagen, Denmark). All sequences were demultiplexed using bcl2fastq v2.20.0.422. Demultiplexing was performed by Macrogen Europe and the National High-Throughput DNA Sequencing Centre.

### Adapter trimming, quality control and mapping

Quality control and adapter removal of demultiplexed DNA was performed using BBMap v38.22 (Bushnell, 2014). Adapters were trimmed with BBduk and sequences were filtered for low quality reads, removing those with average quality less than 20, and further filtered using the following options mink = 11, qtrim = rl, minlen = 50, tbo = T, ktrim = r, and k = 23. Remaining sequences were aligned to the *Fraxinus excelsior* reference genome (BATG0.5) (Sollars *et al*., 2017), and converted to SAM format using BWA v0.7.17-r1188 (Li & Durbin, 2009), and to BAM format using Samtools v1.10 (Li *et al*., 2009). Sequenced reads which did not align to the *F. excelsior* reference genome and read pairs which mapped to mitochondrial and chloroplast regions were removed from the analysis. Remaining sequences that mapped to the nuclear genome gave a mean sequencing depth of 9x with a standard deviation of 8.4 (Supporting Information Table S4).

### Genotype likelihoods, imputation, and genotype probabilities

Genotype likelihoods were calculated for polymorphic positions that passed quality control checks in the *F. excelsior* reference genome (SNP sites) within ANGSD v0.935 (Korneliussen *et al*., 2014). The likelihoods were calculated using the SAMtools model (gl 1) and a minor allele frequency (MAF) of 0.05; sites with data for fewer than 30 individuals were removed (minInd); a minimum base quality score of 20 (minQ) was used; excessive mismatches were removed (-C 50); reads with a flag above 255 (remove_bads 1) and read pairs which did not map correctly were removed (only_proper_pairs 1); reads that had multiple best hits were removed (uniqueOnly 1); finally a *P*-value threshold of 2 x 10^-6^ was used to call SNPs. Furthermore, the BAQ algorithm was included in the analysis to remove false SNP calls due to misalignment of indels (Li, 2011). To reduce computational time ANGSD was launched in parallel, using GNU parallel v20220422 (Tange, 2022). Due to the varying level of coverage between samples of aligned whole genome sequences, missing values were imputed, and subsequently genotype probabilities were calculated in Beagle v3.3.2 (Browning & Browning, 2007). An overview of the computational pipeline is shown in Fig. 1.

**Fig. 1.**
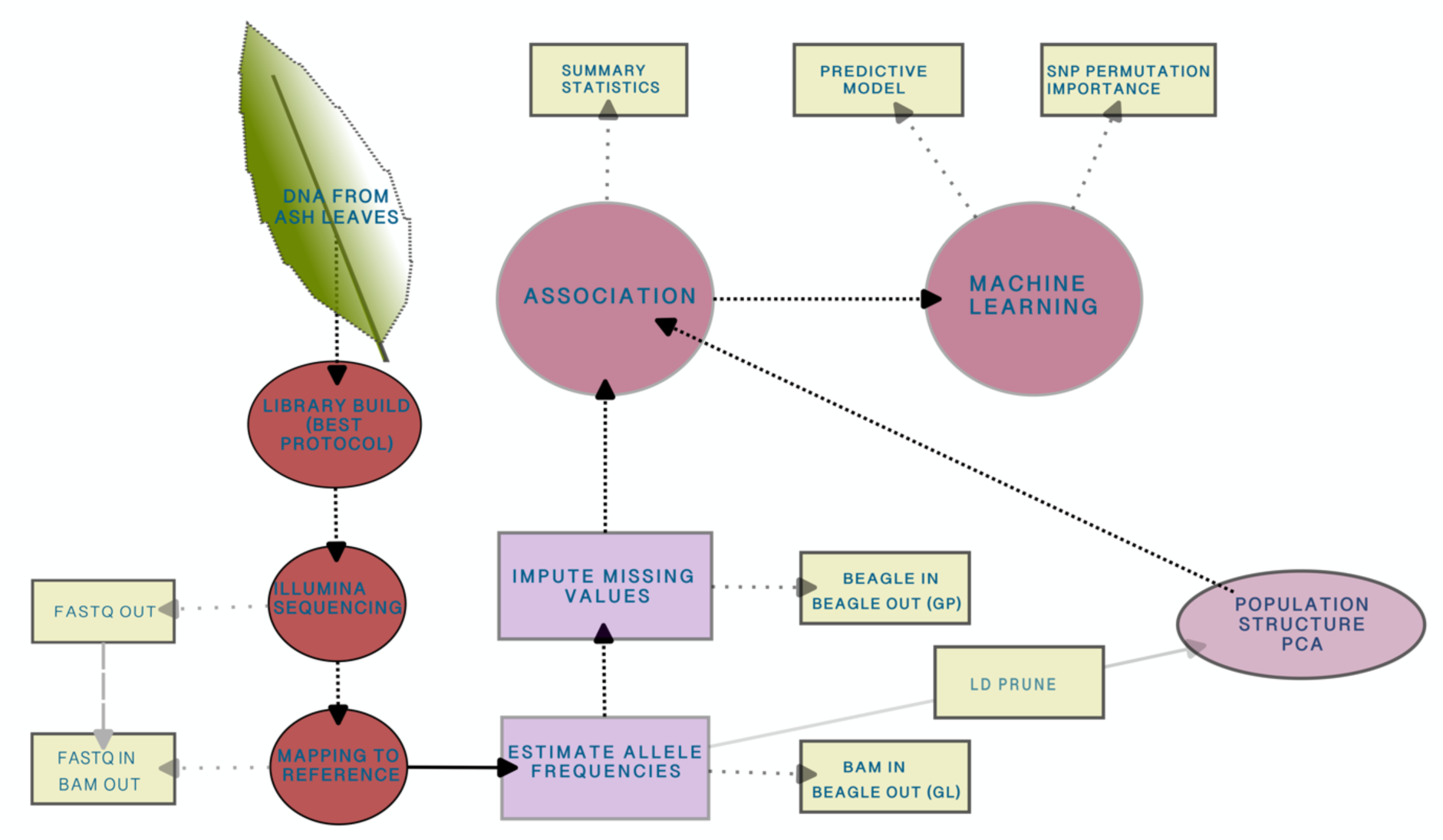
Flow chart of laboratory work and computational pipeline. After collecting *F. excelsior* leaves DNA was extracted from a leaflet, sequenced, and analysed using GWAS and machine learning. GL = genotype likelihoods, GP = genotype probabilities, LD = linkage disequilibrium, PCA = principal component analysis.

### Population structure

Population structure was inferred using SNP genotype likelihoods from ANGSD and assessed using PCAngsd v0.97 (Meisner & Albrechtsen, 2018). Prior to calculation of principal components, linked SNPs were pruned using Plink2 v1.90beta6.24 (Purcell *et al*., 2007), and the indep-pairwise setting with a window size of 50, a variant count of 0.5 and a pairwise R^2^ value of 0.5. As the clones within the clonal *F. excelsior* trials were expected to be unrelated, we expected only low levels of population structuring. However, to prevent confounding factors arising during subsequent association tests, a covariance matrix was estimated and included in subsequent association tests. Eigenvectors were calculated in R from the covariance matrix using the R base function ‘eigen’. Significance values of eigenvectors were calculated using the Tracy-Widom test within the R package AssocTests (Wang *et al*., 2020). There were three significant principal components (eigenvectors) which were included as covariates in association tests. A PCA showing sample relatedness is shown in Supporting Information Fig. S3.

### Association testing

We performed a genome wide association analysis for all quantitative traits using a generalised linear framework implemented in ANGSD v0.935 (Skotte *et al*., 2012). The following options were selected for the association test, ‘doAsso 6’ as the dosage model was used, ‘yQuant’ as phenotypes were quantitative, ‘Pvalue 1’ to print *P*-values, ‘doMaf 4’ to estimate major/minor allele frequency from genotype probabilities, ‘minHigh 30’ to require a minimum of 30 credible genotypes, ‘minCount 30’ to require a minimum of 30 credible minor alleles and the ‘cov’ option which contained the first three covariates from the PCAngsd output. Resultant *P*- values were ranked (the lowest *P*-value SNP was ranked as number 1) and used as input to a machine learning model. SNPs were not corrected for multiple tests. An association for each genotype and phenological trait was created independently.

### Machine learning using Random Forests

The most significant SNPs with the lowest *P*-value from each trait in ANGSD association testing were exported to an R implementation of Random Forests (v4.6-14) for a genotype-phenotype linear regression analysis. Before running the model, SNPs were called and converted to HapMap format. That is, top ranked SNPs based on *P*-value were converted from Beagle format containing genotype probabilities to VCF using FCgene v1.0.7 (Roshyara & Scholz, 2014), then SNPs were called and converted to HapMap format using a custom script available here: https://github.com/clydeandforth/gwas_ash_adapt. SNPs were used as dependent variables, representing the genotype and the phenotypic traits were independent variables represented by a continuous scale. The random forest ensemble model was conducted for each phenotypic trait independently. Within each model, hyperparameter settings such as *mtry* and *ntree* were tested and settings, which provided the most stable model, were used.

Genetically distinct clones were randomly split 50%50% into training (n=243) and test (n=243) datasets. Both the training and test datasets contained the genotypic and phenotypic data for 50% of the samples. The model was trained on the training dataset and then used to predict the phenotypic score of the test dataset. The linear relationship between the predicted score and the actual phenotypic score was evaluated using an adjusted R^2^ value. To understand the ability of varying quantities of SNPs to explain the phenotype (adjusted R^2^), SNPs were divided into groups (cohorts) by lowest *P*-value, i.e. the lowest *P* value was SNP rank 1 and the 10th lowest was SNP rank 10. Based on these rankings, the SNPs were allocated into cohorts of the lowest 10, 50, 100, 500, 1000, 10000, 25000, 50000, 100,000 and 1,000,000. For each cohort a train/test-run was conducted 100 times (using a different seed number for each run, thereby giving a unique split of the data in each run). Conducting the train/test-run 1000 times did not change the results. The mean R^2^ of each permutation set for each cohort was taken as the final adjusted R^2^ value, and the highest adjusted R^2^ value indicated the cohort of SNPs that best predicted the phenotypes (best cohort).

### Functional enrichment analysis

The best performing cohort of SNPs from the machine learning models were aligned to the *F. excelsior* reference genome, and structural annotations from the alignment were extracted. Structural annotations containing SNPs in FASTA amino acid format were used as input to the KOBAS-i online webserver (Bu *et al*., 2021). The organism most closely related to F. excelsior in the KEGG pathway database was *Olea europaea* var. *sylvestris* which was used as a reference for pathway overrepresentation analysis. Significantly overrepresented functional groups were calculated using Fisher’s exact test.

### Permutation importance and feature selection

The predictive ability of each SNP within the maximal cohort for each phenotypic trait (i.e. the number of SNPs which gave the highest R^2^ value) was evaluated using the importance function in Random Forest, by selecting type=1 and scale=FALSE. The value assigned to each SNP as a feature is the permutation importance (PI) (Debeer & Strobl, 2020). The PI is a numeric value which can be positive or negative; a higher PI value indicates higher predictive power. To account for linkage disequilibrium (LD), SNPs were pruned as described above using Plink v1.90beta6.24 (Purcell *et al*., 2007), as LD can considerably affect importance measure calculations (Meng *et al*., 2009). The final SNP sets for each trait used in permutation importance calculations are presented in Supporting Information Data S1-5.

## Results

### Genetic correlation between ash dieback crown damage and phenological traits

Our data shows that estimates of genotypic correlation between ADB crown damage and phenology vary substantially among clonal trial populations, but autumn leaf yellowing was consistently and significantly negatively correlated with crown damage in all trials, except Ireland which showed the same trend but lacked significance (Table 1). This negative correlation means that the more a tree shows leaf yellowing at a given timepoint, the less it is affected by ash dieback. Genotypic correlations for the three remaining phenotypic traits were variable, as autumn status showed a significant negative correlation to ADB crown damage at three of the sites, autumn leaf loss showed a positive correlation to ADB crown damage in two of the trials assessed and spring budburst had both positive and negative correlations to ADB crown damage. Due to the consistent significant negative correlation between ADB crown damage and autumn leaf yellowing, genomic associations for these traits are presented below. Data for the remaining three traits are mainly presented in supplementary information.

**Table 1.**
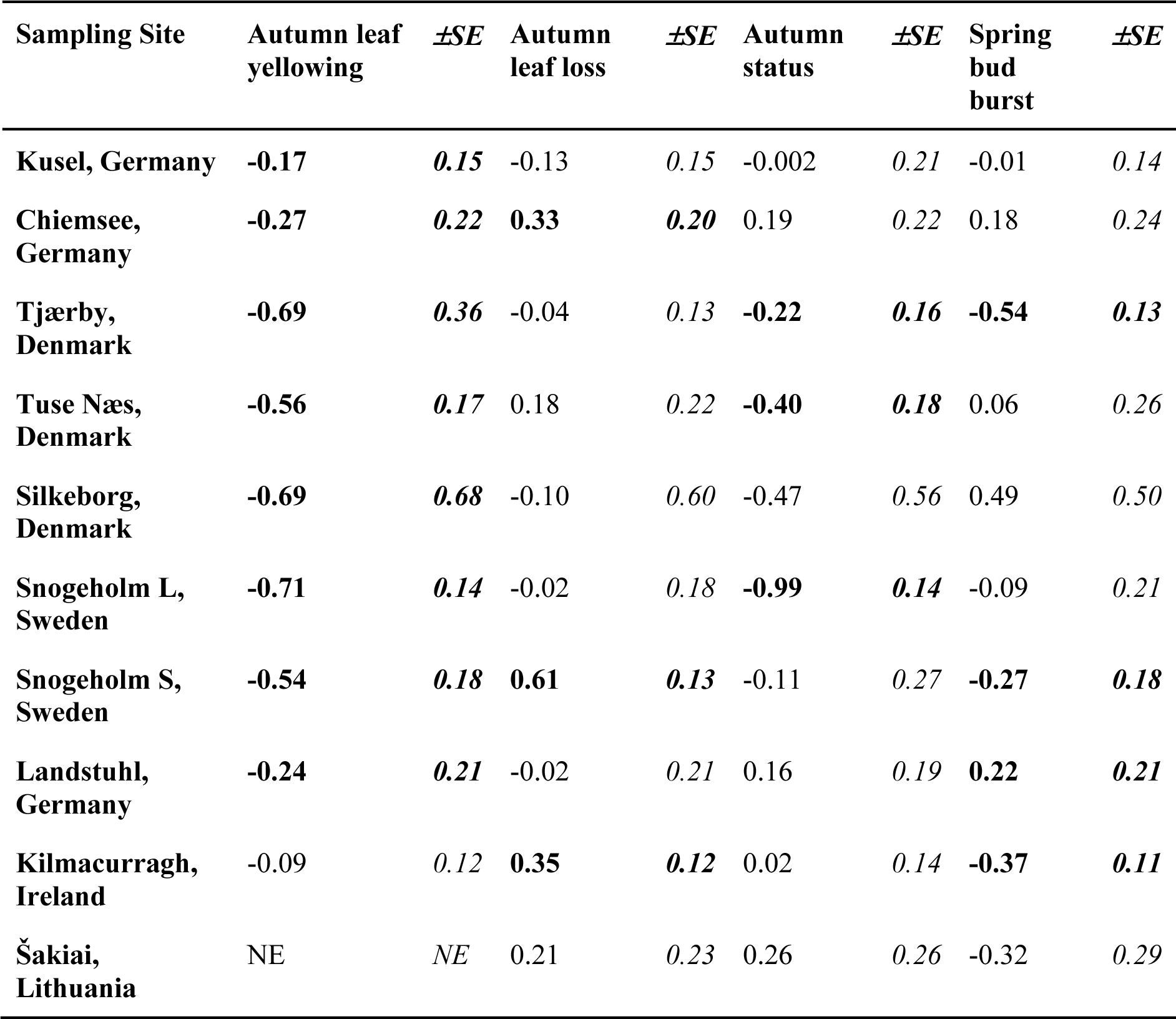
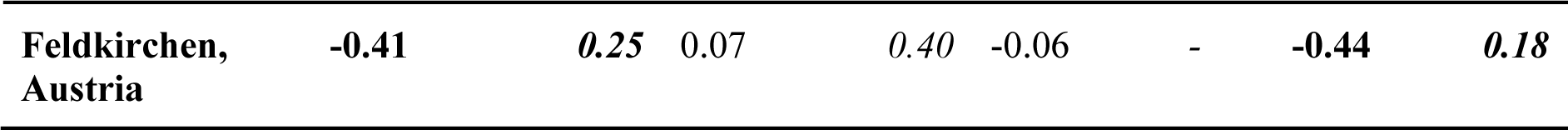
Genotypic correlations between ADB crown damage and phenological traits in *Fraxinus excelsior.* Correlations in bold are significant. SE=standard error. NE=non-estimable due to lack of genetic variation.

| Sampling Site | Autumn leaf yellowing | $\pm SE$ | Autumn leaf loss | $\pm SE$ | Autumn status | $\pm SE$ | Spring bud burst | $\pm SE$ |
| --- | --- | --- | --- | --- | --- | --- | --- | --- |
| Kusel, Germany | **-0.17** | **0.15** | -0.13 | 0.15 | -0.002 | 0.21 | -0.01 | 0.14 |
| Chiemsee, Germany | **-0.27** | **0.22** | **0.33** | **0.20** | 0.19 | 0.22 | 0.18 | 0.24 |
| Tjærby, Denmark | **-0.69** | **0.36** | -0.04 | 0.13 | **-0.22** | **0.16** | **-0.54** | **0.13** |
| Tuse Næs, Denmark | **-0.56** | **0.17** | 0.18 | 0.22 | **-0.40** | **0.18** | 0.06 | 0.26 |
| Silkeborg, Denmark | **-0.69** | **0.68** | -0.10 | 0.60 | -0.47 | 0.56 | 0.49 | 0.50 |
| Snogeholm L, Sweden | **-0.71** | **0.14** | -0.02 | 0.18 | **-0.99** | **0.14** | -0.09 | 0.21 |
| Snogeholm S, Sweden | **-0.54** | **0.18** | **0.61** | **0.13** | -0.11 | 0.27 | **-0.27** | **0.18** |
| Landstuhl, Germany | **-0.24** | **0.21** | -0.02 | 0.21 | 0.16 | 0.19 | **0.22** | **0.21** |
| Kilmacurragh, Ireland | -0.09 | 0.12 | **0.35** | **0.12** | 0.02 | 0.14 | **-0.37** | **0.11** |
| Šakiai, Lithuania | NE | NE | 0.21 | 0.23 | 0.26 | 0.26 | -0.32 | 0.29 |
| Feldkirchen,<br>Austria | -0.41 | 0.25 | 0.07 | 0.40 | -0.06 | - | -0.44 | 0.18 |

### Genotype-phenotype associations reveal signatures of ash dieback and autumn leaf yellowing within genomic linkage blocks

Associations between genomic variants (SNPs) and phenology traits were tested for significance. Across phenological traits SNPs followed an additive model with a polygenic genetic architecture (Supporting Information Figure S4 & S5). This is consistent with previous plant GWAS (Demirjian *et al*., 2023). Linkage blocks displayed using Manhattan plots revealed associated SNPs (Supporting Information Fig. S4 & S5). There were no SNPs which met standard GWAS significance thresholds (5 x 10^-8^ ; Chen *et al*., 2021) in the ash dieback crown damage association, but there were eight SNPs which were significant in the autumn leaf yellowing association. The top five linkage blocks (i.e. those with more than 20 SNPs represented within the 1000 SNPs with the lowest *P*-value) for the ash dieback crown damage phenotype contained no SNPs which aligned to functional regions (i.e. within a genic region or 500 bp upstream or downstream of the genic region; Supporting Information Fig. S4). Similarly, the top five linkage blocks for the autumn leaf yellowing phenotype contained no SNPs within functional regions (Supporting Information Fig. S5). The top ranked SNPs i.e. those with the lowest *P*-values were used as input data for Random Forest linear regression model.

### High phenotypic variance is explained by a relatively low number of SNPs

SNPs were fitted into a Random Forest linear regression model where the maximum adjusted R^2^ value for each trait was calculated by placing SNPs into cohorts between 10 and 1 million based on their GWAS *P*-value, with for example cohort 10 being the SNPs with the 10 lowest *P*-values for that trait association. This approach revealed that the maximal number of explanatory SNPs for observed genotypes (adjusted R^2^) was 25k (adjusted R^2^ = 64%) for ADB crown damage, 100k (adjusted R^2^ = 75%) for spring bud burst, 10k (adjusted R^2^ = 63%) for autumn leaf yellowing, 100k (adjusted R^2^ = 84%) for autumn leaf loss and 50k (adjusted R^2^ = 78%) for autumn status (Fig. 2). High predictive power as indicated by adjusted R^2^ values for each trait were reached using far lower numbers of SNPs. For example, 50 SNPs explain 45% of the genotypic variance for ADB crown damage, increasing to 48% for 100 SNPs to 57% for 5k SNPs before the maximal R^2^ of 64% is reached with 25k SNPs.

**Fig. 2.**
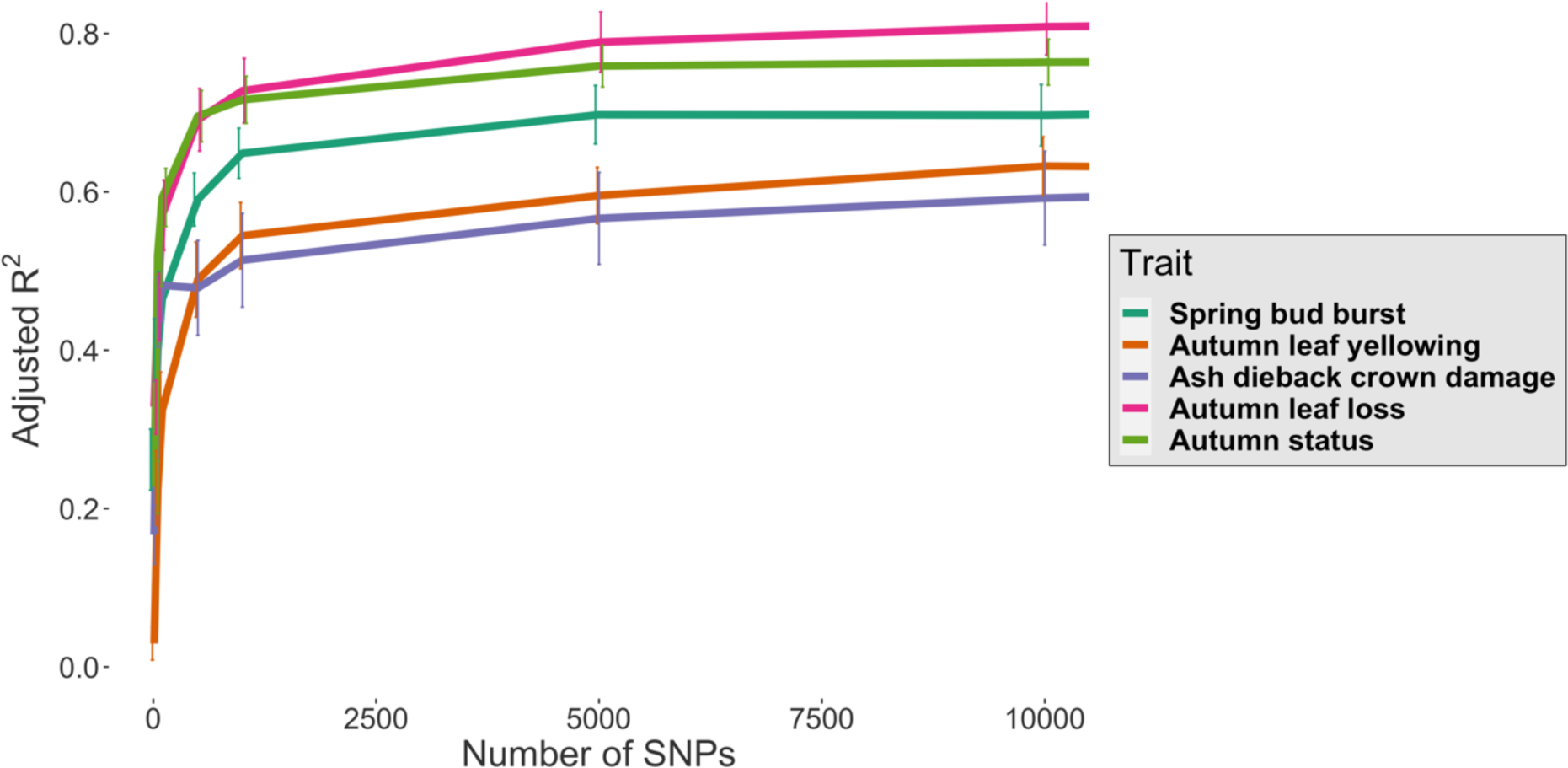
Average adjusted R^2^ for five phenology and ash dieback crown damage traits using the top 10 to 10000 ranked SNPs from Random Forests machine learning models. The observed phenotype for five traits was explained by the explanatory genotype (SNPs) with an adjusted R^2^ value between 0 and1 (displayed in the y-axis).

Similarly, the pattern of variance explained in Fig. 2 for the autumn leaf yellowing trait shows a more gradual rise to reach the plateau, as the first 10 SNPs have an adjusted R^2^ value of 3% which increases to 32% with 100 SNPs and 54% with 1k SNPs before reaching a peak of 63% at 10k SNPs. The remaining three traits (spring bud burst, autumn leaf loss and autumn status) required more SNPs to reach their maximal adjusted R^2^ value at 50k (autumn status) and 100k (spring bud burst and autumn leaf loss) SNPs, and as with the ADB crown damage trait have a rapid increase in proportion of variance explained from relatively few SNPs (Fig. 2). Overall, there was shows a general trend of increasing SNP number explaining a greater proportion of phenotypic variance for all traits (Fig. 2).

The cohort of SNPs resulting in the highest adjusted R^2^ value was used for functional enrichment and permutation importance (PI) analysis. Permutation importance is a measure of model performance where each feature (here SNPs) is randomly shuffled, and the error rate of the model calculated (Nicholls *et al*., 2020). For example, as 25k SNPs produced the maximal adjusted R^2^ value for ADB crown damage, SNPs from this cohort were first assessed for enrichment of the gene function based on their loci. Secondly, the SNP set was pruned for linkage disequilibrium and the reduced number of SNPs were thereafter ranked according to their permutation importance. Selected SNPs were those, that were top ranked from a machine learning model of PI (i.e. having the greatest effect on phenotype prediction). SNPs which remained after pruning were the final list of SNPs for each trait and those which were used for further analysis (Supporting Information Data S1-S5).

### SNPs associated with ADB crown damage are located within coding regions of genes related to stress tolerance and phenology

Ash dieback crown damage is an overall measure of the damage inflicted on individual trees by *H. fraxineus*. Associations between ADB crown damage and SNP variants gave the highest explanatory effect using 25k SNPs (adjusted R^2^=63%). After pruning for linkage disequilibrium, a final list of 9155 SNPs remained. These SNPs were ranked based on their PI score. The loss of explanatory information after removing linked SNPs was negligible as the 9155 SNPs still had the same value (when digits were rounded), with an adjusted R^2^ of 63%. The linked SNPs which were pruned from the dataset had similar predictive ability with a marginally reduced R^2^ of 62%, revealing redundancy among the 25k SNPs which is likely due to linkage disequilibrium.

Alignment of the 25k SNPs to gene loci on the ash reference genome and a subsequent clustering analysis, provided a profile of enriched functional groups associated with ADB crown damage (Fig. 3). Among the enriched functional groups were biosynthesis of secondary metabolites and sesquiterpenoid and triterpenoid biosynthesis. These enriched defence-related functional groups indicate coding variation in tolerance and phenology genes, which may reflect the observed phenotypic variation in tolerance. For example, the biosynthesis of secondary metabolites group includes phytoalexins and phenolics, known antimicrobial, communication and stress-response compounds (Bhattacharya *et al*., 2010; Ahuja *et al*., 2012). This indicates a variable defence response among *F. excelsior* individuals to *H. fraxineus*.

**Fig. 3.**
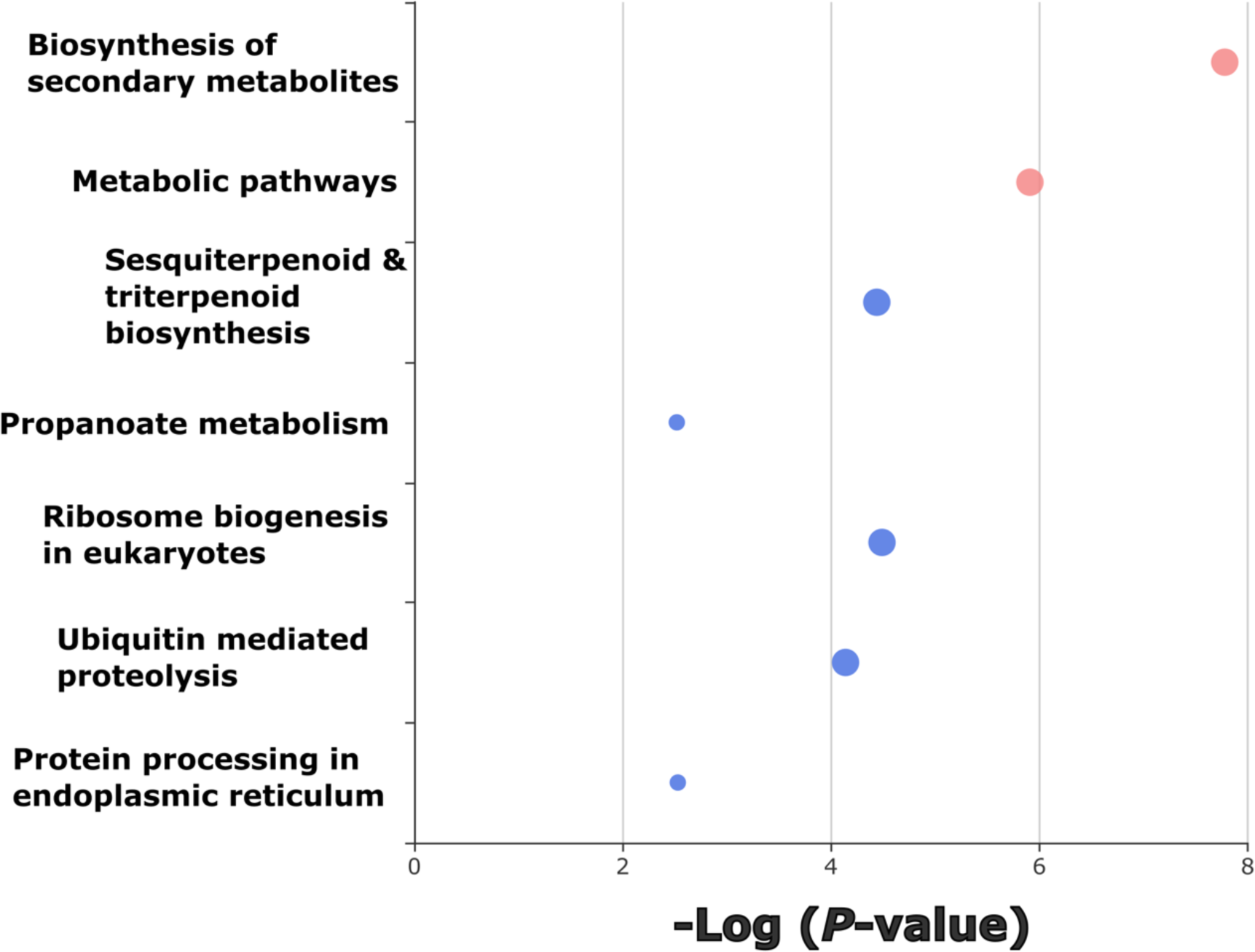
Enrichment analysis of genomic regions comprising the 25k single nucleotide polymorphisms (SNPs) with the highest explanatory power within a machine learning model for ash dieback crown damage. Colours represent related functional groups (in this figure only pink/red circles represent related functional groups), as pathways were calculated using a network algorithm (Rosvall & Bergstrom, 2008). Note that blue circles indicate unrelated functional groups. Bubble size represents the magnitude of enrichment for each functional group. Displayed bubbles have a *P*-value <=0.05.

The PI score and genomic location (i.e. within a coding domain, intragenic region, up or downstream of a gene, intergenic region or untranslated region) of the top 100 ranked SNPs is shown in Fig. 4. SNPs within the top 100 for PI score which were located on or near to genes were functionally annotated using the NCBI and KEGG databases. Among the top 100 SNPs, three were on coding domains; all three SNPs cause non-synonymous, missense mutations which change the translated amino acid (Table 2). In addition three SNPs were located in untranslated regions (SNP 2, telomerase reverse transcriptase; SNP 6, presenilin At1g08700; SNP 56 disease resistance RPM1-like) and although these are not translated into proteins they have important effects on transcriptional regulation, such as mRNA stability and translation which have demonstrable effects in human disease (Hindorff *et al*., 2009; Steri *et al*., 2018; Conteduca *et al*., 2021). Of the remaining top 100 SNPs, seven were located up or downstream of a gene, 13 were within intragenic regions (introns), and the remaining 74 were within intergenic regions.

**Fig. 4.**
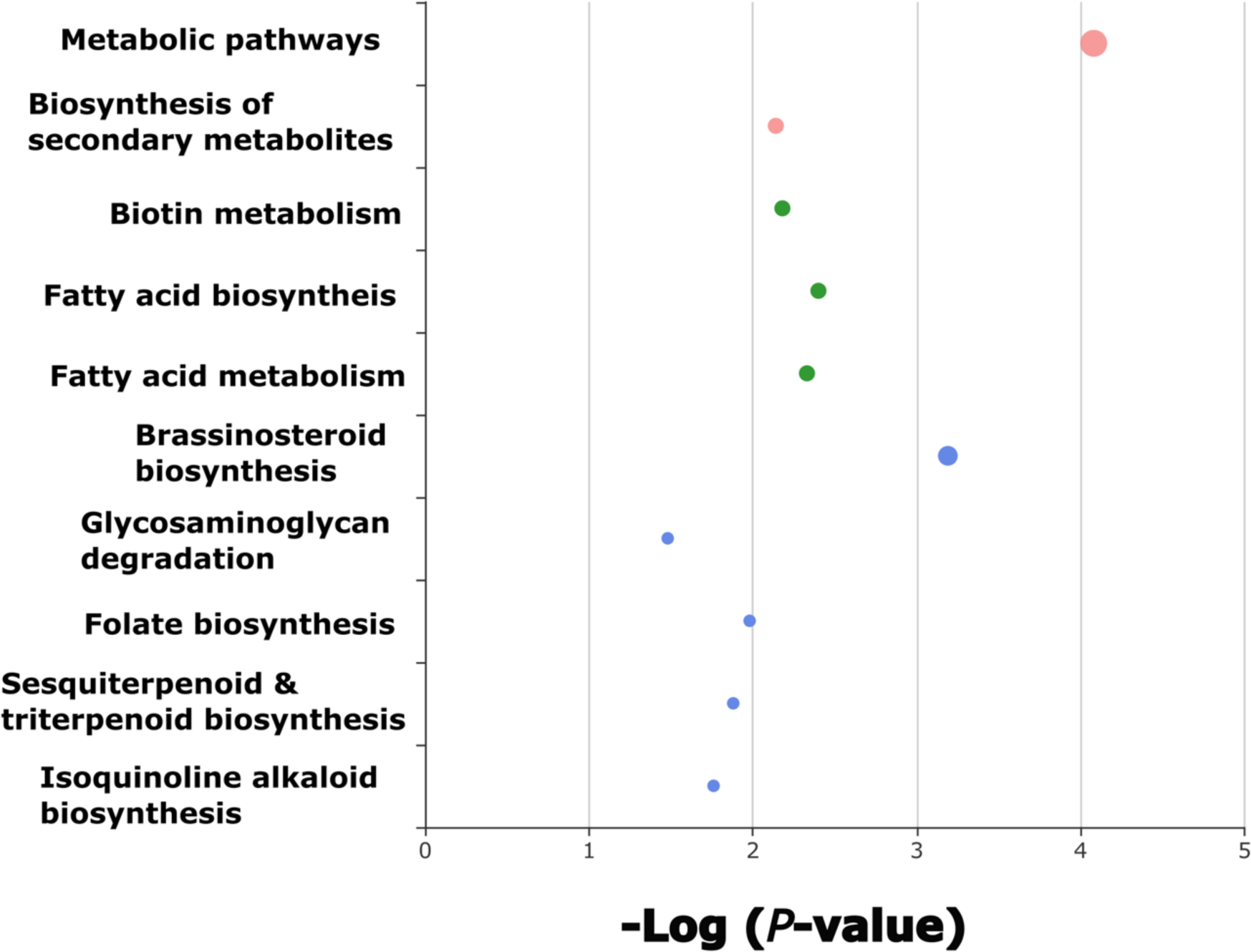
Top 100 SNPs ranked by permutation importance associated with ash dieback crown damage in *F. excelsior* individuals. SNPs were ranked from a panel of 9155 SNPs, ranked according to their permutation importance. SNPs are marked according to their position within a genic region: SNPs within 500 bp of genic regions are marked ‘Up/Downstream’; SNPs resulting in a coding domain substitution (CDS) are shown in green and the value given in the y-axis is their PI rank with GWAS rank in parentheses. SNPs which cause missense mutations have functionally annotated coding domains. Permutation importance score is shown in the x- axis.

**Table 2.**
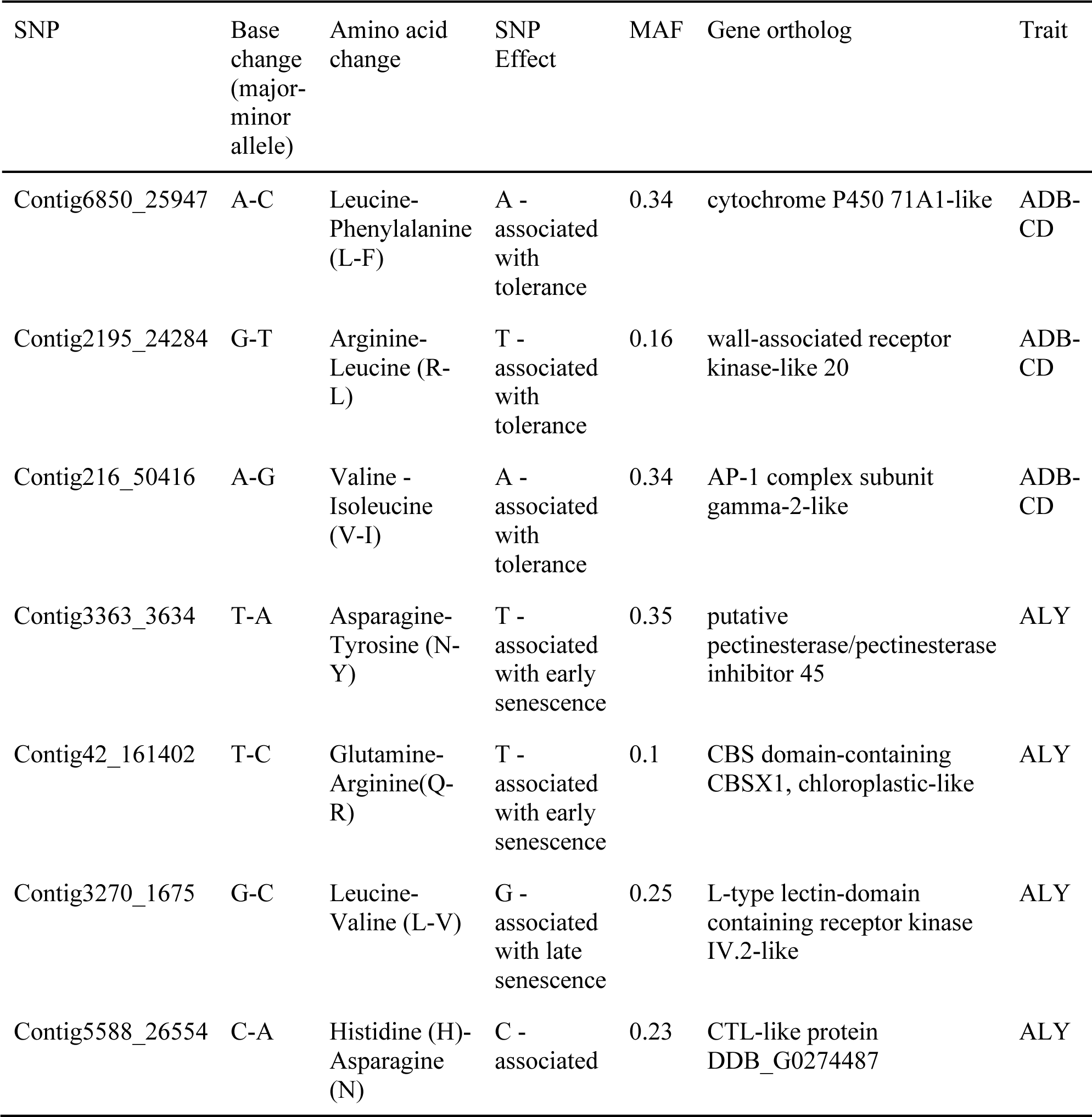
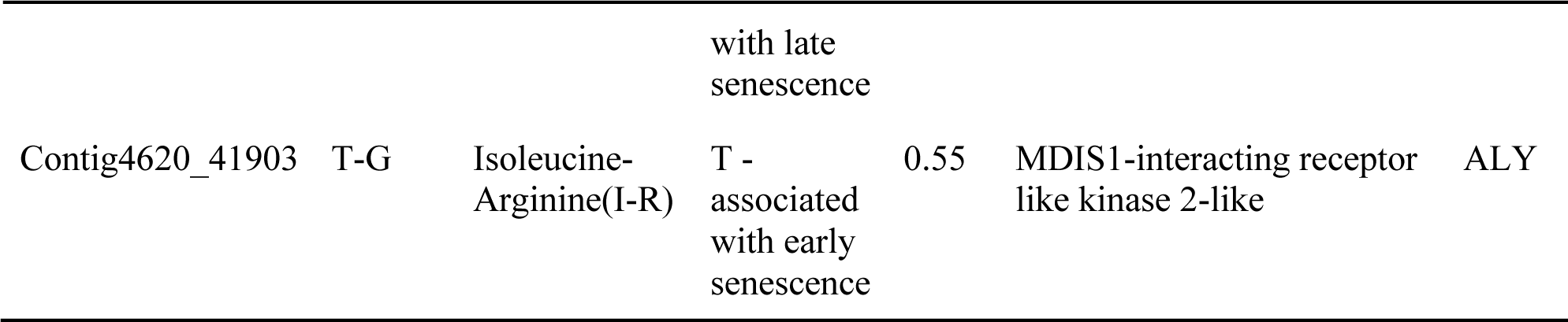
List of non-synonymous SNPs with high permutation importance, located within functionally important genes. ADB-CD = ash dieback crown damage. ALY = Autumn leaf yellowing

The highest ranked SNP within a coding domain was the 57th ranked SNP overall by PI and by GWAS ranking had the 17961st lowest *P*-value (Fig. 4). This SNP was located within a gene region coding for a cytochrome P450-71A1-like protein which is involved in avocado fruit ripening (Bozak *et al*., 1990), but in which a mutation on a gene homolog in barley suppresses the development of necrotrophic fungal pathogens through reinitiating programmed cell death, a plant defence mechanism that reduces access to nutrient rich plant tissues and which can be hijacked by pathogen elicitors (Ameen *et al*., 2021). The second of the three SNPs is located within a region encoding a wall-associated receptor kinase-like 20 protein; homologs of this gene are part of the pattern triggered immunity (PTI) system, a conserved plant defence system which recognises conserved components of plant pathogens and triggers an immune response (Jones & Dangl, 2006). The third and final coding domain SNP was located on an AP-1 complex subunit gamma-2-like gene which in *Arabidopsis* is critical for the development of male and female gametophytes (Zhou *et al*., 2022).

### Autumn leaf yellowing - pathogen triggered immunity and phenological regulation are associated with trait variation

Autumn leaf yellowing is a quantitative measure of loss of green pigment in leaves. Colour change in *F. excelsior* is usually green to yellow, but may stay green before falling in central Europe (Roloff & Pietzarka, 1997). Genotypic prediction of autumn yellowing based on the top 10k ranked associated SNPs could explain 63% of this trait. Post LD pruning there were 3740 SNPs remaining. This increased the R^2^ to 72% suggesting that some pruned SNPs did not add to the explanatory power but instead decreased the accuracy of the model and did not contribute to an accurate genotype prediction. However, the removed linked SNPs (*n*=6260) gave a R^2^ of 58% suggesting that many SNPs were linked and like the ADB crown damage genotype reveals redundancy in the larger (10k SNP) dataset.

Enrichment analysis based on the 10k SNP dataset reveals functional groups related to plant defences such as brassinosteroid biosynthesis, sesquiterpenoid and triterpenoid biosynthesis and isoquinoline alkaloid biosynthesis (Fig. 5). Given that autumn leaf yellowing is correlated to ADB crown damage, SNPs within these functional groups may be a component of ash tolerance to *H. fraxineus*.

**Fig. 5.**
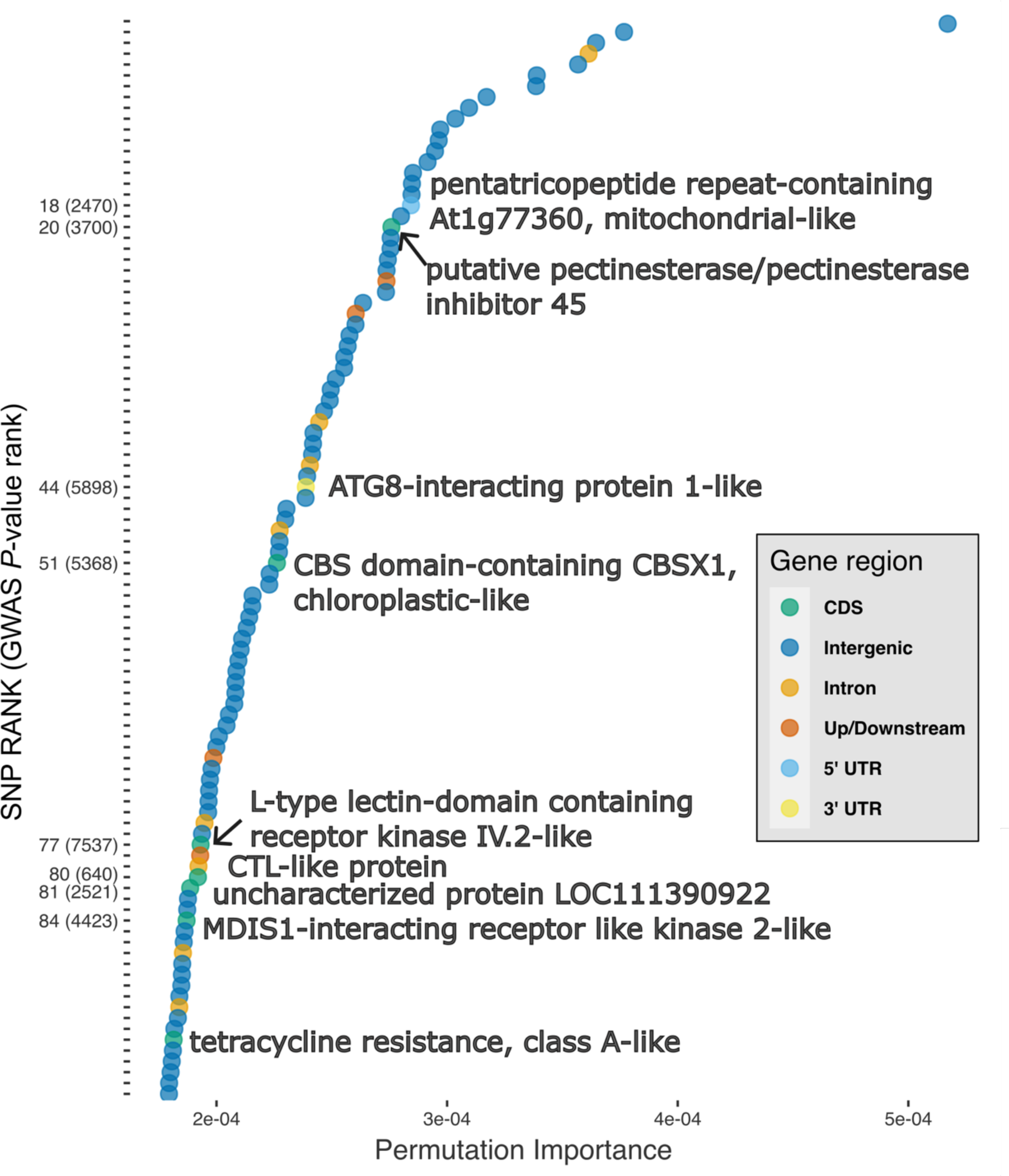
Enrichment analysis of genomic regions comprising the 10k single nucleotide polymorphisms (SNPs) with highest explanatory power within the machine learning model for autumn leaf yellowing. Colours represent related functional groups, as pathways were calculated using a network algorithm (Rosvall & Bergstrom, 2008). Note that blue circles indicate unrelated functional groups. Bubble size represents the magnitude of enrichment for each functional group. Displayed bubbles have a *P*-value <=0.05.

Within the top 100 SNPs with the highest PI, seven were within coding domains, (where six produced non-synonymous amino acid substitutions), two were in untranslated regions, four were up or downstream of a gene, eight were within introns and the remaining 79 were on intergenic regions (Fig. 6). Of the seven SNPs on CDS, the first at SNP rank 20, was a putative pectinesterase with a pectin methylesterase inhibitor domain on the 5’ region of the gene. Pectinesterase inhibitors have known anti-virulence functions as they mitigate the effect of pectin lyase effectors, which are a common component of the virulence arsenal in many plant pathogens (Dean *et al*., 2012; Liu *et al*., 2018). Two genes with non-synonymous SNPs were located within gene regions for fungal elicitor recognition and are homologous to components of the PTI system (Jones & Dangl, 2006). As previously described, PTI is an evolutionary conserved pathway which recognises molecular patterns and upon recognition activates defence components such as the oxidative burst. Specifically these are annotated as an *L*-type lectin-domain containing receptor kinase IV.2-like, ranked at SNP 77 (Wang *et al*., 2014, 2018) and an MDIS1-interacting receptor like kinase 2-like ranked at SNP 84 (Coleman *et al*., 2021). The penultimate plant defence related SNP in a coding domain was ranked at 81 and has no characterised protein homologs, and the final SNP, ranked at 95 encodes a synonymous nucleotide on a coding domain (tetracycline resistance, class A-like). There were two SNPs causing non-synonymous amino acid substitutions on gene coding domains which have functional homologs related to phenology regulation. The first of these at SNP rank 54 is within both a 3’UTR and a coding domain, which was functionally annotated as a homolog of CBSX protein which regulates flowering and leaf maturation through the thioredoxin system. The second amino acid substitution is within a choline transporter like-1(CTL-1) which transports auxin. Auxin is a plant hormone, which regulates many plant development functions including leaf abscission and yellowing (Jin *et al*., 2015).

**Fig. 6.**
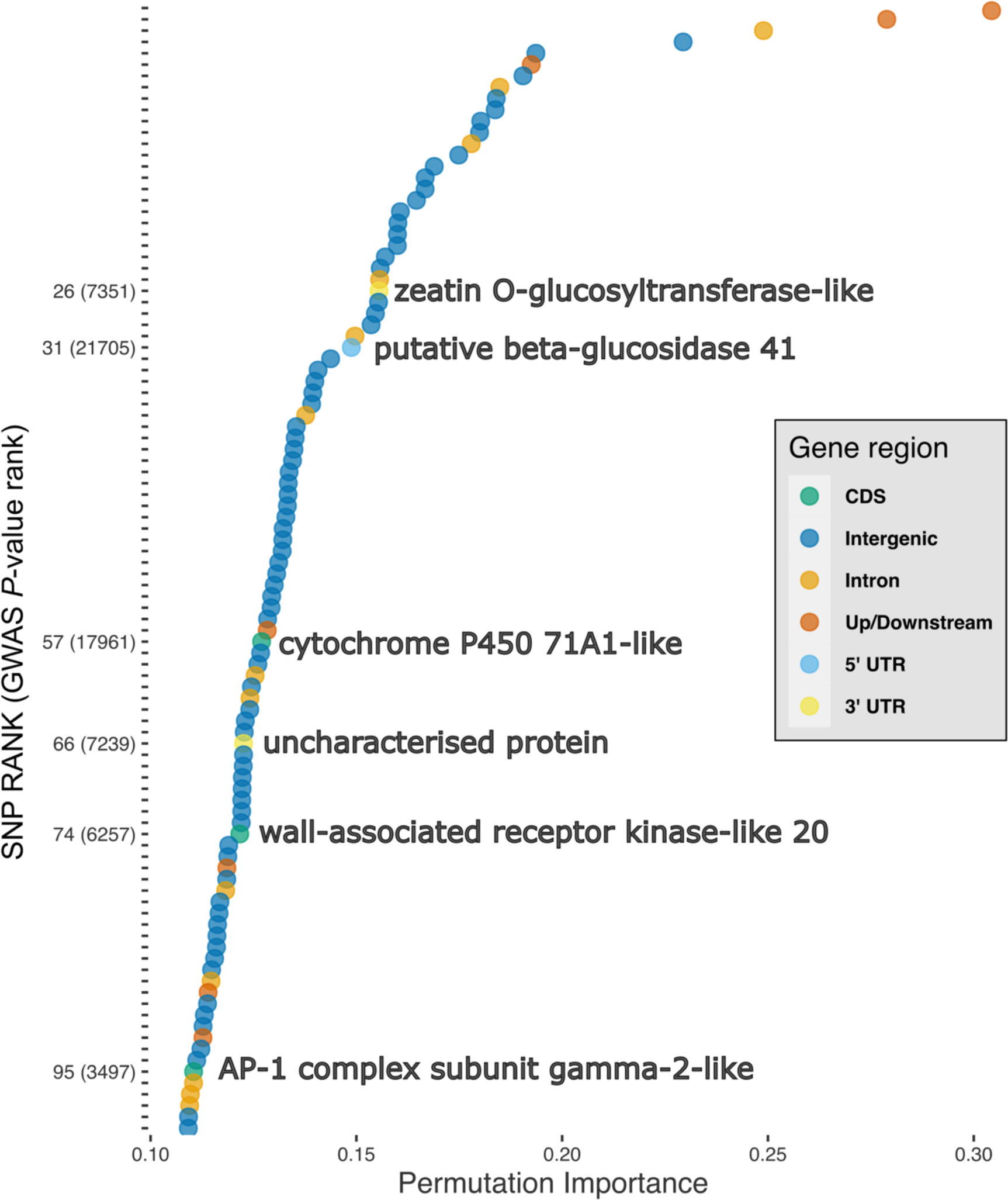
Top 100 SNPs associated with autumn leaf yellowing in *F. excelsior* ranked according to their permutation importance (PI) from a panel of 3740 SNPs. SNPs are marked according to their position within a genic region, SNPs within 500 bp of genic regions are marked ‘Up/Downstream’, SNPs resulting in a coding domain substitution (CDS) are shown in green and the value given in the y-axis is their PI rank with GWAS rank in parentheses. SNPs which cause missense mutations have functionally annotated coding domains. Permutation importance score is shown in the x-axis.

## Discussion

Results from previous common garden studies suggest that susceptibility to ADB is under polygenic control with moderate to high genetic narrow-sense heritability (*hns* =0.3-0.5) (Pliura *et al*., 2011; Kjær *et al*., 2012; Lobo *et al*., 2014; Muñoz *et al*., 2016). Here, we found that associated SNPs could collectively explain a similar fraction of the phenotypes as we would expect based on heritability estimates (e.g. up to 63% based on 9155 unlinked SNPs for crown damage score). Our findings suggest that ash dieback susceptibility is based on many genes with small effects in line with GWAS results from Stocks *et al*. (2019). Furthermore, we found no overlap between genes associated with crown damage level and autumn leaf yellowing - two genetically correlated traits. Although, this is potentially confounded by the impact of ash dieback on progression towards senescence. The results are in support of the observation that genetic variation in susceptibility is likely based on many rather than few genes with additive contribution towards susceptibility. Our findings are important in relation to the likely dynamic of future host-pathogen interaction, as tolerance controlled by a single gene or a few genes is more likely to be broken down compared to tolerance based on many genes (Ennos, 2015). Despite the relatively low number of SNPs accounting for a large proportion of susceptibility, it would take many evolutionary steps for the pathogen to overcome multiple tolerance associated SNPs, each with a small effect on the phenotype. Surpassing the tolerance offered by a single SNP would not make a large quantitative difference to the overall tree phenotype. Combined with recent results which present evidence that the biology of the pathogen does not favour selection of high virulence (Kosawang *et al*., 2020), and evidence of superior reproductive success of less susceptible individuals, gives some hope for the future of common ash (Semizer-Cuming *et al*., 2019, 2021). High mortality among the current generation is an acute concern but higher fitness of tolerant individuals, combined with restoration efforts and breeding programs will strongly support the continuation of *F. excelsior* in European forests.

Our findings are based on genotypes from several European trials where the genotypes have been carefully classified as superior or inferior (in terms of susceptibility to ADB), based on long-term monitoring. We used a common protocol in data collection for the present study; trees with different genetic backgrounds were evaluated at different ages and sizes, and in trials with different growth conditions. The outcome of the genome-wide SNP analysis is, thus, an accurate prediction of ash dieback susceptibility and phenological traits across pan Central-North European *F. excelsior* populations. It presents an important step towards a better understanding of the genetic control of ash tree susceptibility in populations exposed to this emerging pathogen. The results are therefore relevant, reproducible, and representative of the ash population across the entire disease outbreak area.

### Genomic background of correlation between autumn leaf yellowing and ADB susceptibility

Our results provide new insight into the intriguing strong genetic correlation previously reported from common garden studies, where early leaf yellowing was associated with lower levels of susceptibility to ADB (McKinney *et al*., 2011; Stener, 2013). By assessing phenology and ADB susceptibility simultaneously, we confirmed the genotypic correlation between autumn leaf yellowing and ADB susceptibility assessed as crown damage score (Table 1). Moreover, we compared SNPs associated with each of the two correlated traits and found that a relatively small number of SNPs can also explain the genetic control in autumn leaf yellowing (e.g. adjusted R^2^ of 33% with the 100 top ranked SNPs). Functional analysis of these SNPs revealed their enrichment of plant defence related genes (Fig. 5). Associated SNPs did not overlap (i.e. the SNPs were not the same) in the 100 top ranked SNPs in the correlated traits of ADB crown damage and autumn leaf yellowing. However, analysis of the molecular functions of the SNP loci suggested possible pleiotropic effects as for example defence components were associated with autumn leaf yellowing (Table 2).

### Functional analysis of genes in correlated traits

We searched for SNP variants associated to correlated traits that offer indications on how trees avoid severe disease, i.e. SNPs within functional regions which code for phenology and defence related genes. Within the autumn leaf yellowing trait, SNP variants code for amino acid substitutions in the regulation of flowering and leaf abscission (CBSX-1), and an auxin transporter (CTL1-like protein). These genes have homologs in plants which alter the timing of seasonal growth cycles (Wang *et al*., 2017; Murai *et al*., 2021). It is possible that the substitutions in these amino acids reflect observed variation in early change in autumn leaf colour as first reported by McKinney *et al*., 2011. Genetic correlation between phenotypic traits can either be a result of essentially the same genes controlling the traits (pleiotropy) or as a result of linkage disequilibrium (Lynch & Walsh, 1998); in the latter case it must be explained by the evolutionary history or a bias created during selection of the tested genotypes.

Correlations between ADB crown damage and autumn leaf yellowing, together with associated SNPs support the asynchronous growth theory, i.e. trees avoid severe disease by early leaf abscission (McKinney *et al*., 2011; Harper *et al*., 2016). We did, however, not find a significant genetic correlation between ADB crown damage and autumn leaf loss, but as leaf loss is also a symptom of infection by *H. fraxineus*, leaf yellowing may be a more robust measure of autumn senescence, including shedding of leaves. The high level of genotypic correlation was based on genotypes selected before they were challenged by the novel pathogen (McKinney *et al*., 2011), and therefore selection bias is not likely to be an important explanation. Early shedding of leaves may create some level of disease escape, but our results suggest that tree defences are the major tolerance factor (Figs. 3-6 & Table 2). The data indicated that the two traits were controlled by different genes but still presented high genetic correlation. This suggests that genetic control of susceptibility is due to host defences and a possible coincidental relationship to autumn leaf yellowing. However, if the variation in susceptibility among individuals was caused only by autumn leaf yellowing, then the top ranked SNPs would be the same in ADB crown damage and autumn leaf yellowing. The view that active defence against the spread of the pathogen in host tissue is involved is also supported by the results from inoculations with *H. fraxineus* directly onto branches, where genotypes with low susceptibility to natural infections also showed limited bark necrosis development; furthermore development of necrotic bark lesions occurred much faster than in genotypes with high susceptibility (McKinney *et al*., 2012), suggesting that an active defence against the spread of the pathogen is involved. As there is no co-evolutionary history between *H. fraxineus* and common ash it is likely to be a broad-spectrum defence mechanism which limits the impact of *H. fraxineus* tolerant ash genotypes.

### SNPs in PRR related genes are associated with tolerance to ADB

Our results further show that standing genetic variation (i.e. the available allelic variation within a population at a given timepoint (Barrett & Schluter, 2008)) within tree defences is associated with variation in disease severity across the European ash population (Figure 2). This raises the possibility that individual variation in tolerance to ADB is mediated by host recognition of molecular patterns within *H. fraxineus*. Pattern recognition receptors (PRRs) are components of the plant defence system, and intriguingly, four of the six PRR related genes identified from top ranked SNPs are associated with autumn leaf yellowing. These PRR related genes are known to stimulate plant defence components (Boutrot & Zipfel, 2017). As leaf yellowing is correlated with ADB crown damage, it may be evidence of gene pleiotropy (Table 2). As previously suggested, an active defence is likely to be a key component of tolerance to ADB, and recognition of pathogen components of the hemibiotroph *H. fraxineus* by *F. excelsior*, is a possible mechanism of tolerance to ADB (Lobo *et al*., 2015; Mansfield *et al*., 2018; Nielsen *et al*. 2017). Here, we provide putative genomic evidence of ADB tolerance among individuals through pattern triggered immunity mediated recognition of an external threat and a response to that threat through defence activation.

## Conclusions

This study highlights the genetic architecture and possible mechanisms involved in the genetic control over ADB crown damage and phenology traits. The implications of these findings are: 1) ADB tolerance is polygenic, but a large proportion of the variance can be explained by a relatively small number of SNPs (Fig. 2); 2) At the gene level, there is evidence of disease tolerance via plant defence recognition of *H. fraxineus*. Furthermore, phenological avoidance of severe disease is a component of tolerance, especially as it is correlated with ADB crown damage. However, a previous study showed that direct inoculation with *H. fraxineus* and the resultant defence response is responsible for individual variation to the pathogen (Lobo *et al*., 2015). Our findings highlight that ADB susceptibility is determined by combinations of polygenic traits. When assessing crown damage levels, we see the combined effect of different mechanisms, i.e. disease tolerance (host trees limiting the growth of the pathogen) and disease escape (autumn leaf yellowing, as a proxy for senescence) but possibly also other mechanisms. A tree that is not able to limit the growth of the pathogen (i.e. showing low tolerance) might still be able to escape the disease due to early senescence (as indicated by autumn leaf yellowing) or perhaps pathogen growth is inhibited in yellow leaves, and hence might show limited crown damage. This emphasises the difficulty in identifying relevant genes in GWAS using crown damage levels or tree health as phenotypic traits. In the future, a better understanding of the molecular defence mechanisms in ash trees in response to the pathogen are needed. One way to approach this might be to target more specific traits, e.g. necrosis length after controlled inoculations might provide a better estimate of disease tolerance which is not confounded with disease escape. However, at the end the complex interplay of different mechanisms determines the survival of ash trees.

Coming generations of *F. excelsior* will rely on a reduced number of ash trees that manage to survive in the face of *H. fraxineus* (an omnipresent sexually recombining pathogen). To ensure the future of *F. excelsior* in Europe, ash trees with a wide range of tolerant phenotypes must be protected and natural regeneration should be fostered to maintain a reservoir of genetic diversity in the European population. Broad genetic diversity would not only protect the species from *H. fraxineus* but also from native and incoming threats such as global change and imminently, the invasive Emerald Ash Borer approaching Central Europe from Russia and Ukraine.

## Data availability

Supplementary data is available here: https://github.com/clydeandforth/gwas_ash_adapt/tree/master/Supplementary_data

## Acknowledgements

We thank Lene H. Andersen, Jacob A. Rasmussen and Sarah Mak for help with lab work, and Lars N. Hansen, Darius Kavaliauskas for assistance with ash trial assessments and Michael Schott for assistance with sampling.

This research was supported by the Independent Research Fund Denmark (DFF|Technology and Production Sciences) under the grant no. 8022-00355B. BML DaFNE grant no. 101476 (“Esche in Not - Phase II”) funded by theAustrian Federal Ministry of Agriculture, Forestry, Regions and Water, Management, the Austrian Chamber of Agriculture, the governments of all Austrian provinces and “Österreichischer Forstverein”. BML DaFNE grant no. 101684 (“Waldfonds” project “AshBack”) funded by the Austrian Federal Ministry of Agriculture, Forestry, Regions and Water Management.

## Notes

https://github.com/clydeandforth/gwas_ash_adapt/tree/master/Supplementary_data

### Competing Interest Statement

The authors have declared no competing interest.

